# Localist neural plasticity identified by mutual information

**DOI:** 10.1101/658153

**Authors:** Gabriele Scheler, Martin L Schumann, Johann Schumann

## Abstract

We present a model of pattern memory and retrieval with novel, technically useful and biologically realistic properties. Specifically, we enter n variations of k pattern classes (n*k patterns) onto a cortex-like balanced inhibitory-excitatory network with heterogeneous neurons, and let the pattern spread within the recurrent network. We show that we can identify high mutual-information (MI) neurons as major information-bearing elements within each pattern representation. We employ a simple one-shot adaptive (learning) process focusing on high MI neurons and inhibition. Such ‘localist plasticity’ has high efficiency, because it requires only few adaptations for each pattern. Specifically, we store k=10 patterns of size s=400 in a 1000/1200 neuron network. We stimulate high MI neurons and in this way recall patterns, such that the whole network represents this pattern. We assess the quality of the representation (a) before learning, when entering the pattern into a naive network, (b) after learning, on the adapted network, and (c) after recall by stimulation. The recalled patterns could be easily recognized by a trained classifier. The recalled pattern ‘unfolds’ over the recurrent network with high similarity to the original input pattern. We discuss the distribution of neuron properties in the network, and find that an initial Gaussian distribution changes into a more heavy-tailed, lognormal distribution during the adaptation process. The remarkable result is that we are able to achieve reliable pattern recall by stimulating only high information neurons. This work provides a biologically-inspired model of cortical memory and may have interesting technical applications.

## 1 Introduction

Storing patterns and achieving sequence-independent recall is a problem for neural network models. We are interested in exploring the concept of localist ensemble memory in cortical networks (Carpenter, 2000; Fuster, 1998). From biology we know that there are both synaptic and intrinsic plasticity (Langille & Brown, 2018), which have a ‘hidden’ component in cell-internal memory (Scheler, 2023). Here we want to explore the concept of storing memory by local plasticity in neurons within ensemble-like neuronal groups. For this purpose we define a realistic excitatory-inhibitory network with recurrent interactions and use simple, visually-defined patterns (LeCun & Cortes, 2005) as inputs to the network. We then examine the representations that develop on a naive network, where ‘naive’ means without previously stored patterns, and in the absence of plasticity. Representations are classified by a downstream machine learning mechanism. The machine learning mechanism stands for other interpretations, for instance, contralateral, or subcortical brain areas, which are not explicitly modelled (cf. Fig. 1).

**Fig 1.**
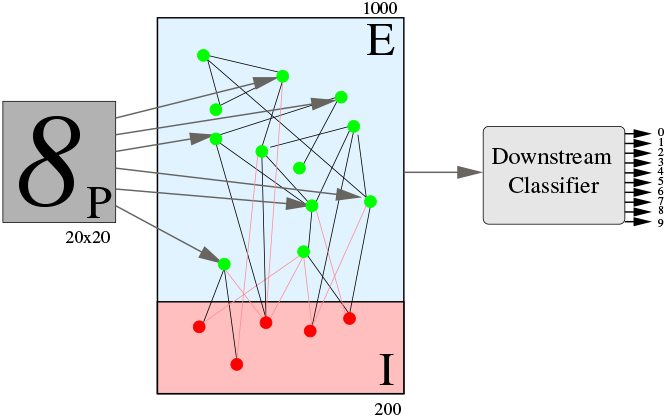
Architecture of the system: patterns from a field of 20×20 excitatory neurons are ‘loaded’ onto E neurons from an E-I network. The network of E (N=1000) and I neurons (N=200) generates a representation. The representation is identified by a downstream (e.g., subcortical) classifier.

The construction chosen is a suitable model for cortical networks (Kim, Leahy, & Shlizerman, 2019; Scheler, 2018), it is also reminiscent of LSN (Maass, Natschläger, & Markram, 2002) and ESN (Jaeger, 2002) models. To implement plasticity, we perform an information-theoretic analysis over neurons and patterns (Schneidman, Bialek, & Berry, 2003; Sheintuch, Rubin, & Ziv, 2022) to find the neurons with the highest mutual information (MI) for each class of patterns. Such high MI response neurons form spontaneously in the representations. The existence of high MI neurons is an emergent property of the network setup. It turns out that while absolute MI values for neurons differ across classes, and the number of high MI neurons above a threshold is different for each class, we can rank the highest MI neurons for each pattern class. We select only the high MI neurons to use for plasticity. The idea is to ‘compress’ the pattern information into the high MI neurons’ intrinsic properties and adjacency network, i.e. perform localist plasticity. Then by stimulating only these neurons, we can recall neural representations and we achieve high precision pattern recall (Scheler, 2016).

We also look at inhibitory neurons which play an important role in defining ensembles in real cortical networks. When an ensemble is activated, inhibitory plasticity guarantees that the activated ensemble is less inhibited, while inhibition remains strong for all other neurons.

When there is low or no overlap in the high MI neurons for each pattern class, localist learning means that there is a disjunctive set of adaptations for each pattern set. Dissimilar pattern classes are stored free of interference. The goal of our experiments is to ensure that pattern information can be retrieved by stimulation of a few, high MI neurons alone - even though the patterns were learned from a full input representation.

## 2 Methods

### 2.1 Neurons and Network Structure

The network model is initialized as a fully connected model with E (=1000) excitatory neurons and I (=200) inhibitory neurons, plus P (=400) pattern input neurons, which are directly linked to 400 of E excitatory neurons, in order to ‘load’ patterns of length P into the network. The connections between the P neurons and corresponding E neurons is strong enough to ensure solid transfer of activity. Synaptic connections within the network are of type NMDA, AMPA and GABA-A and are modeled as in (Scheler, 2018).

Neurons are modeled as spiking neural models as in (Izhikevich, 2004) with an equation for the membrane model *v* and an equation for the gating variable *u* (Eq 1), such that *v* is set back to a low membrane potential *v* := *c*; and the gating variable *u* is increased by an amount *d* (*u* := *u*+*d*), when a neuron spikes (at *v*(*t*) = *θ* and *θ* = 30*mV*).

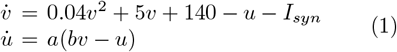

For excitatory neurons, parameters are variable in order to capture different types of neurons and generate different distributions of intrinsic excitability. For excitatory neurons, parameters a and b are varied in our model (cf. (Scheler, 2018)), which results in different intrinsic excitability of a neuron, cf. Table 1. The gain *g* is captured by *g* = *γ* · *a*. We measure the baseline rates as spontaneous neural activity with a background input of around 0.15*nA*. This variation of parameters also allows for intrinsic plasticity.

**Table 1.** Parameter ranges for excitatory and inhibitory neurons.

| Parameter | E | I | P |
| --- | --- | --- | --- |
| $a$ | 0.006 ... 0.034 | 0.3 | 0.1 |
| $b$ | 0.06 ... 0.34 | 0.1 | 0.1 |
| $c$ | -70 | -70 | -70 |
| $d$ | 3 | 2 | 3 |
| $\theta$ | 30mV | 30mV | 30mV |
| $\gamma$ | $2.5 \cdot 10^4$ | $5.0 \cdot 10^4$ | $2.0 \cdot 10^4$ |
| *baseline rate (Hz at 0.15nA)* | 3-7 | 20 | 6 |

For inhibitory neurons, the parameters are

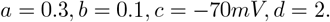

This results in inhibitory neurons firing around 20Hz. For the input pattern neurons P, the values are similar with a lower gain and longer delay (*a* = 0.1, *b* = 0.1, *c* = − 70*mV, d* = 3).

We initialize excitatory neurons with a Gaussian distribution over its gain and baseline rate, such that we use a mean of *µ* = 5 and distribution *σ*^2^ = 0.96 for the rate. Attested values for cortical neurons in mice are rate distributions with *µ* = 4.96 and *σ*^2^ = 0.31 (Scheler, 2017).

Initially, the network has full connectivity for E-E, E-I and I-E synapses. There are no I-I connections. Synaptic strength is set to achieve baseline activity in the network with a background input (∼ 0.15*nA*). Accordingly, the network is initialized with a Gaussian distribution of synaptic strength for both AMPA and NMDA connections between E-E (*µ* = 0.0015, *σ* = 0.00027) and E-I (*µ* = 0.014, *σ* = 0.00024). For GABA-A connections, we use a Gaussian distribution with *µ* = − 0.5 and *σ*^2^ = 0.1.

There is a signal delay between neurons, which is randomly distributed:

- AMPA: 10 … 30*ms*
- NMDA^1^: 10 … 15*ms*
- GABA-A: 5 … 15*ms*

We do not use this parameter during plasticity.

### 2.2 Patterns

In order to use a simple set of variations of patterns which can be easily classified into classes, we used the MNIST database (LeCun & Cortes, 2005) for 10 handwritten digits, with 50 variations for each digit. The format was an integer vector (for the grayscale values) of length 400 (10×50=500 patterns).

We load patterns into the E+I (=1200) processing network via P (=400) pattern neurons which have a single excitatory connection (AMPA) to 400 E neurons in the processing network. We apply one input pattern of length 400 at a time for 300ms. We measure the resulting neural representation for the pattern after end of application. Spike rates recorded from excitatory neurons (E=1000 neurons) for a length of about 300 ms are regarded as the neural representation of the pattern. After that time, the neural representation fades away in our set-up.

### 2.3 Classification of Representations

In our work we wanted to give an overall impression of whether the adaptation method used could demonstrate pattern learning (i.e. correct interpretation of new, never-seen patterns). We used an ML system, AutoGluon (Erickson et al., 2020; Gijsbers et al., 2023; Schumann, 2025), which is a supervised training method using a mixture-of-experts approach. It employs several mechanisms for pattern classification in parallel - all of them have access to the pattern representation, and they are tested against a validation set. It achieves its performance by combining models from a variety of learning algorithms such as Random Forests, K-Neighbors, Categorical Boosting, Gradient Boosting Machines, Neural Networks, and combined ensemble models (Erickson et al., 2020). The results are accumulated and those with the better prediction are weighted stronger for the goal of pattern classification of new patterns.

After training the basic models, the predictions of these models on a validation data set are used to train a meta-model, which will yield a weighted combination of predictions of the basic models. Figure 2 shows how three basic learning algorithms are combined to achieve a combined classification result, which has a score value higher than each of the basic components.

**Fig 2.**
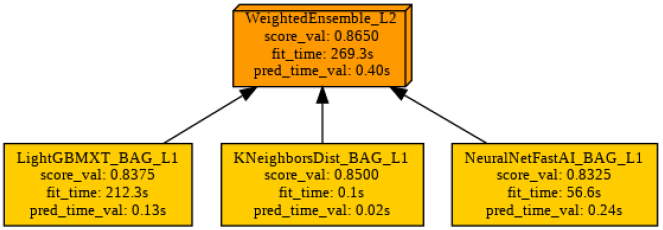
AutoGluon: combination of classification models in a stack. The “score val” is the score for the validation data set. The weighted combination (Model Weighted Ensemble L2) achieves a higher value by a learnt weighted combination of the predictions of the 3 individual models.

We wanted to use this to demonstrate that acceptable pattern learning occurs. We made no attempt of improving our results, or of tackling complex datasets.

The Autogluon classifier is presented with the neural representations for the 50 * 10 = 500 patterns. The classifier’s task is to recognize the correct digit for each pattern. The original neural representations for patterns are labeled, and the algorithm is trained to classify them (‘supervised learning’).

Combining different pattern classification methods is very likely also a technique that is used in biology to arrive at best results, results which may be best suited to specific tasks. In this respect it is remarkable that cortical activation patterns are accessible to (one could say “observed by”) different networks: contralateral cortical networks, striatal networks, hippocampus, midbrain areas like substantia nigra and VTA, the cerebellum etc. Quite obviously the ability to classify patterns is different in all those brain areas, and this diversity serves many purposes, such as task-dependent classification results, and will have different levels of precision. There is no need to assume that there is a single learning algorithm used by the brain.

The test phase consists in recognizing representations from stimulating of a few high MI neurons *alone* after neural plasticity. These ‘recalled’ neural representations were derived from stimulating *m* high MI neurons where *m* = 5, 10, 20. We can show that these representations are recognized by the classifier based on their similarity with the original representations. This is a remarkable result.

### 2.4 Information Theoretical Analysis

Each pattern class *s* is presented with equal probability *p*(*s*) = 1*/N*_*digits*_, where *N*_*digits*_ = 10.

The response for each neuron is the rate response *r*. To simplify, we distinguish 3 rate responses (A,B,C), with 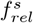 as the neuron’s firing rate for digit pattern *s* relative to its baseline rate (Eq. 2).

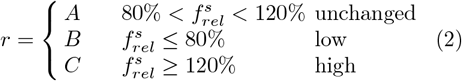

Now for each set of 50 variations of a digit, each neuron has a probability for each of the three responses. For instance, for digit 5, neuron j may have *A* = 0.8, *B* = 0.1, *C* = 0.1, or 40 times A, 5 times B and 5 times C. Assume that overall *p*(*A*) = 0.5, *p*(*B*) = 0.25, *p*(*C*) = 0.25.

We calculate the mutual information (MI) (Zeng, 2015) between each digit *s* (10 with 50 variations) and rate response *r* for each neuron *n* as in (Eq. 3):

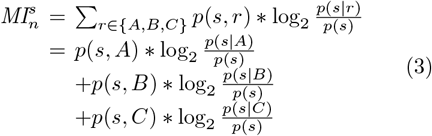

Assume *p*(*digit*5) = 0.1 overall. Since A is the ‘neutral’ option of a fluctuating value in an intermediate range, neuron j would not be very specific in recognizing pattern 5. Its mutual information 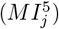 would be (0.054 + (− 0.01322) + (− 0.01322) = 0.0278 *bit*).

Figure 3 shows the overall MI for each excitatory neuron of the network after processing 500 patterns without any plasticity. The neurons are arranged according to their intrinsic baseline firing rate (right-to-left, and top-to-bottom). MI is higher after plasticity. There is no correlation with the firing rate.

**Fig 3.**
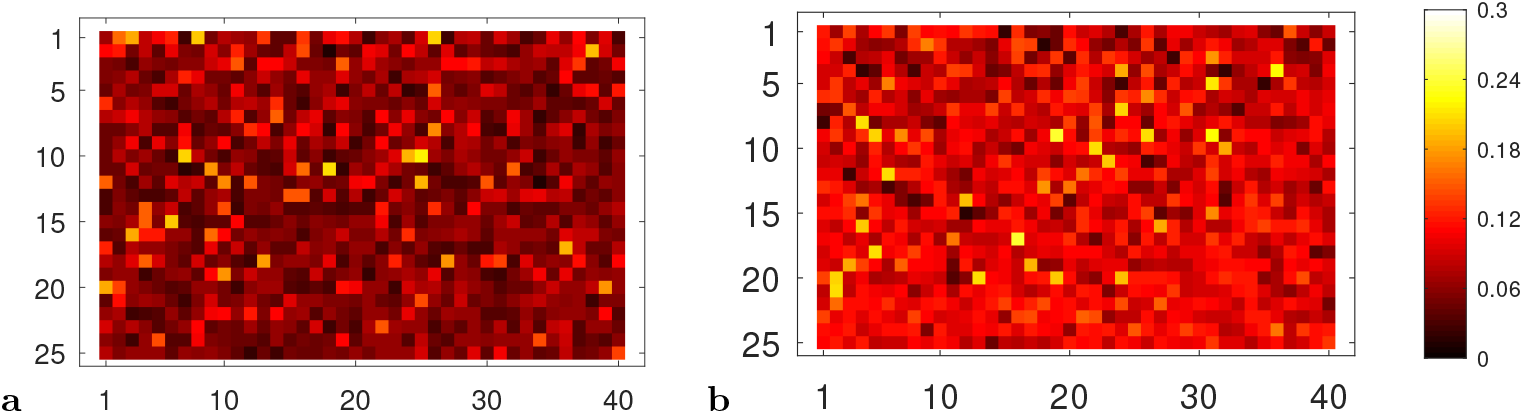
Increase of MI by plasticity. No correlation between baseline firing rate and MI. 1000 E neurons are shown on a 25×40 grid. The numbering is ordered by the baseline firing rate of each neuron (left to right and top to bottom). MI is calculated for all 500 patterns. (**a**) before plasticity is applied, (**b**) after plasticity is applied.

We can also calculate the mutual information for each neuron with respect to each digit pattern. For each neuron, we have 10 values for 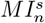, since there are *s* = 1..10 digit patterns. We then rank the *m* highest 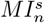 values for each digit pattern. The result is shown in Fig. 4**a**–**c**. There are no shared high MI neurons for *m* = 5, and few for *m* = 10, *m* = 20.

**Fig 4.**
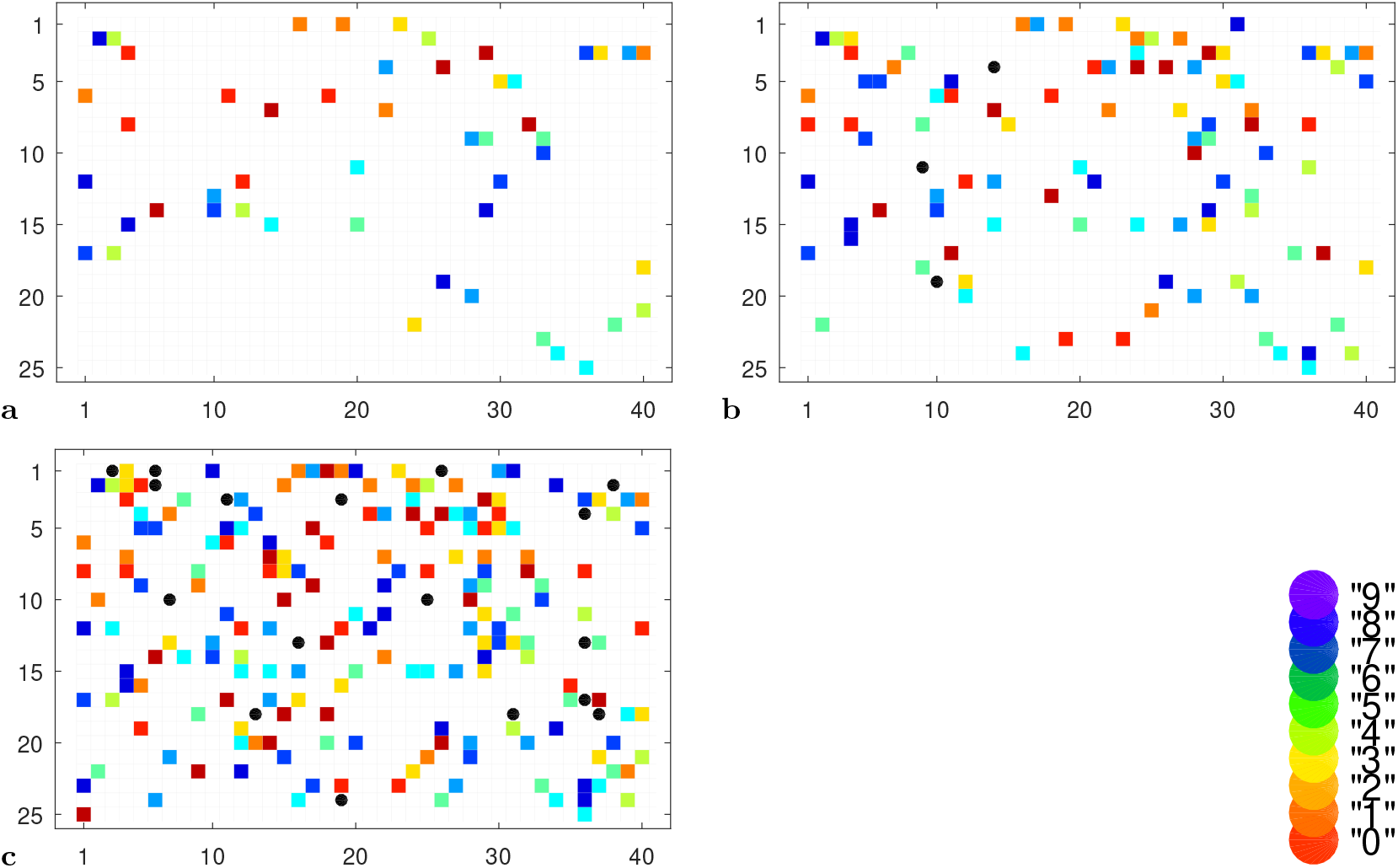
Ranked high MI neurons (*m*) for a naive network representation, unique for each digit (0-9), marked by color. (**a**) *m* = 5 neurons per pattern, (**b**) *m* = 10 neurons, and (**c**) *m* = 20 neurons. Shared high MI neurons are marked as round and black. There are no shared high MI neurons for *m* = 5. They are sparse even at higher *m*.

### 2.5 Plasticity

Adaptation is based on a neuron’s information content (MI). The system has access to the mutual information of a neuron for patterns. A neuron must have high *MI* (for some pattern) in order to undergo plasticity. For each pattern, we rank neurons by mutual information, and select the *m* highest neurons which are active for this pattern, for *m* = 5, 10, 20 (Fig. 4). Surprisingly, the specific size of *m* shows little difference for the behavior of the system.

We use two types of single-shot update rules, for intrinsic plasticity, and for synaptic plasticity.

- Intrinsic plasticity: For the adaptation of high MI neuronal parameters *a, b*, we use an exponential update rule:

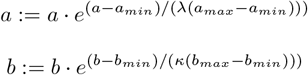

With this rule, we strengthen the excitability in a non-linear fashion with stretch factors *λ* = 3 and *κ* = 3. It turned out that a proportional update rule (*a*· (1 + *λ*)) did not lead to desired results.

When parameters *a* and *b* were already high, and the neuron had a high intrinsic activation, further adaptation of the parameters could cause undesirable behavior, such as intense bursting-like activity. This is an artifact of using a parameterized neuron model which is non-optimal for extreme values. Lowering the learning rate overall would lead to a lack of intrinsic learning for the bulk of low-firing neurons. Therefore we chose an exponential form, which ensured that learning intensity was matched to the existing activity level: strong learning for low activity neurons, weaker learning for high activity neurons. It is easy to imagine a biological basis for this.

- Synaptic connections:

- For high MI neurons *m* only, their postsynaptic (input) connections are updated according to the activation of the input neurons *i*: if the response activation is high (type C, Eq 2), the connection will be strengthened

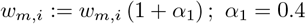

This parameter *α*_1_ was set manually such that recall was optimized. For responses A (unchanged) and B (low), *w*_*m*,*i*_ is left unchanged. A single update of synaptic value is sufficient.

- Presynaptic (output) connections are updated according to the activation response of the neurons *o*: if the response is high (type C, Eq 2), the presynaptic connection is strengthened:

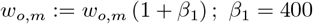

Here *β*_1_ ∼ 400 shifts the weight to a much higher value.

Synaptic connections for output neurons *ω* with unchanged or low responses are additionally weakened:

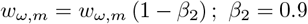

The updates for weakened synapses were not crucial in our experiments, they mainly serve to better calibrate the overall distributions.

- Tuning inhibition:

- For each pattern, a few inhibitory neurons *inh* are selected to receive strengthened connections from pattern-specific high MI neurons *n*:

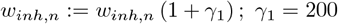
- The GABA-A connections from inhibitory neurons *inh* to excitatory neurons *e* with unchanged or negative response are also strengthened in order to suppress neurons which do not participate in coding for the pattern:

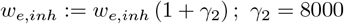

For inhibition, parameters selectively strengthen and suppress synapses and in this way a trace of the neural representation is ‘carved’ into the network.

In this case, meta-parameters were hand-tuned to achieve appropriate results for localist recall. The different magnitudes of the meta-parameters *α*_1_, *β*_1_, *β*_2_, *γ*_1_, *γ*_2_ are explained by the size of the neuronal sets that are being linked, e.g., *α*_1_ for the connections from ∼ 200 type C neurons to ∼ 10 m neurons and *β*_1_ for the connections from ∼ 10 m neurons to ∼ 200 type C neurons. Automatic setting and re-calibration of parameters could be added and might uncover more parameter combinations with acceptable solutions (‘regimes’). It is most likely that calibration by homeostatic plasticity, which occurs from time to time, would also stabilize the parameter regimes, and make some weakening rules unnecessary.

The intrinsic plasticity rule means that a neuron which carries much information for a pattern will increase its intrinsic frequency. It will also adjust its input and output connections. The pattern will leave a trace at that neuron. We have seen that intrinsic frequency and high MI are initially not correlated (Fig. 3). For small sets of high MI neurons, there is limited overlap in the identity of the high MI ‘concept’ neurons for each pattern (Fig. 4), and therefore the update rules are mostly disjoint (separate). A high MI neuron’s input synapses are updated only when they receive positive pattern input from input neurons *i*. However, all of its output synapses are updated. As a result, we have few, local updates which furthermore are applied only once, which makes plasticity highly efficient.

## 3 Results

### 3.1 Neural Representations and Classification

We modeled the network similar to a cortical network, which receives pattern information from thalamic input neurons (Sampathkumar, Miller-Hansen, Sherman, & Kasthuri, 2021). Accordingly, we used a vector representation of visual patterns for perceptual input and added weak fluctuating background noise (of 1.5*nA*). We initialized the network with random (Gaussian) distributions of neural parameters such that the network had heterogeneous intrinsic strengths and synaptic weights. The naive network, without any plasticity, developed neural representations for, in this case, 500 pictures from 10 classes. The neural representations, which lasted ∼ 300*ms*, could be analyzed for information content of individual neurons using the tool of mutual information. It turned out, like we noticed before (Scheler, 2016), that neurons with high mutual information develop for each input pattern class. These neurons could represent pattern classes in the manner of symbolic or ‘concept’ neurons. They stand for the whole pattern. We developed a localist form of plasticity to make pattern representation permanent. Then, by stimulating only very few (*m* = 5 − 10) high-MI neurons, we recall pattern representations sufficiently to allow for correct pattern classification by a machine learning classifier (Section 3.4).

For the naive representations, without any plasticity applied, we show that the projection of the 400 pixel input pattern onto the 1200 recurrent cortical network model resulted in recognizable neural representations. Figure 5 shows neural representations for digit “6”.

**Fig 5.**
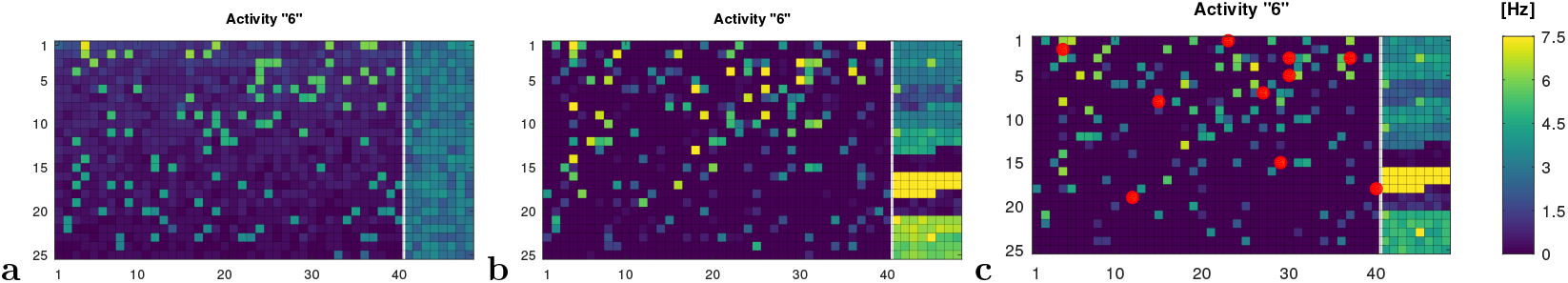
Neural rate representation for digit patterns. Excitatory neurons on the left side of each panel, inhibitory neurons on the right. (**a**) Activity after presentation with a pattern “6” before plasticity, (**b**) The same after application of plasticity rules. (**c**) Activity after stimulation of 10 high MI neurons (red). Patterns (**a**–**c**) were all classified correctly as representing digit “6”. The high inhibitory activity contributes to specificity of representation.

1. before application of plasticity, when the pattern is presented to the naive network through input neurons P (Figure 5**a**),
2. after plasticity, when pattern is presented to the learned network through input neurons P (Figure 5**b**), and
3. after plasticity, when the representation is ‘recalled’ from stimulation of the ‘concept’ high-MI neurons alone (Figure 5**c**).

The results from two different example runs are summarized in Table 2. A reduction of recall precision after adaptation (Figure 5**b**) which allows to generalize to unseen patterns is in accord with many results from the computational ML literature.

**Table 2.** Accuracy of representations during learning. Two different examples (run A and B) are shown.

| run | naive | trained | recall $m = 5$ | recall $m = 10$ | recall $m = 20$ |
| --- | --- | --- | --- | --- | --- |
| A | 472/500 (94%) | 422/500 (84%) | 10/10 | 10/10 | 9/10 |
| B | 477/500 (95%) | 437/500 (87%) | 10/10 | 9/10 | 8/10 |

Figure 8 shows that the number of strong response neurons is reduced after training. Presumably, increased inaccuracy is a result of compression of pattern representation (see below Section 3.4).

### 3.2 Mutual Information Analysis of Representations

When we analyzed the neural representations for information content, by measuring mutual information (MI) between each neuron and the 10 digit input patterns, we could see that there is no or weak correlation between high MI and high baseline rate overall in the network (Figure 3). The correlation is low before plasticity (*r* = − 0.09 (Pearson), − 0.10 (Spearman)) and remains low with *r* = 0.048, 0.06 after plasticity.

After plasticity, the mutual information (for 500 patterns) in the network is higher for each pattern (Figure 6). This shows that the network has stored pattern-specific information. It has increased its information content specifically for each of the learned patterns.

**Fig 6.**
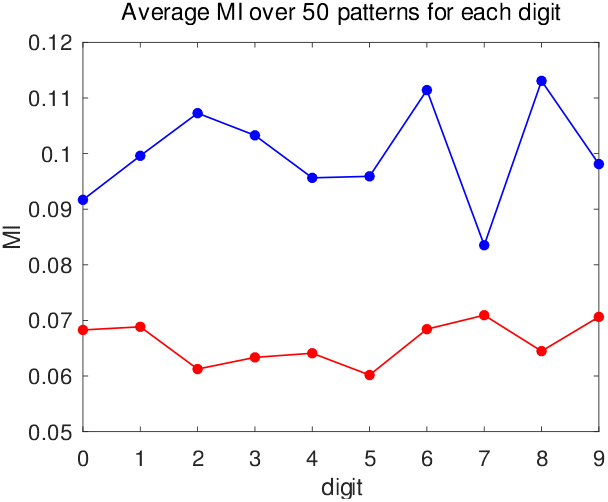
Average MI per digit before applying plasticity (red) and after training (blue) for run A.

Before training we set up a network with different initializations. This is why we get slightly different results in different runs (cf. Table 2). After training the results are fully deterministic (the background input is always the same). The high MI sets give the same results for each recall experiment. The MI system has no internal source of noise, randomness, or indeterminacy.

Our aim was to identify high MI neurons for each digit which could be regarded as ‘symbolic’ abstractions of the whole pattern. I.e., we were looking for the *m* neurons for each digit pattern, which are highest ranked by mutual information. Our results show that these neurons have a low amount of overlap (Figure 4). Most high MI neurons are unique and spread connections over the whole network. No calculations are offered here on the relations between data sets, network size, and number of MI neurons. This would allow to generalize the results of this form of plasticity beyond the chosen example.

### 3.3 Localist Plasticity and Recall

In order to store the representations of pattern input that appear on the naive, randomly initialized network, we apply plasticity rules. Our goal is to compress the information into selected neurons and their surrounding ensembles, and recall patterns by stimulating those high-level, ‘concept’ neurons. There are experimental indications for such constructions (Josselyn & Tonegawa, 2020; Park, Ko, Frankland, & Josselyn, 2024), and they also have enormous advantages in intelligence applications.

We have shown that MI is distributed in such a way that few neurons have high MI. We want to use these neurons like symbolic abstractions. In a number of experimental publications (Agetsuma et al., 2023; Gal et al., 2021), a hub-spoke representation for each pattern has been suggested (Gu, Qi, & Gong, 2019). We believe that such structures imply great advantages for recall. It should be sufficient to target only the high-order neurons for activation, which then activate their feature structures, and in this way reconstruct the whole pattern (cf. also (Choucry, Nomoto, & Inokuchi, 2024; Uytiepo et al., 2024)).

Accordingly, we use a simple, one-shot plasticity rule, focused on high MI neurons. We may label this type of plasticity rule “localist”, because it only affects few neurons and their synaptic environment. The basic idea is to form a trace of the input pattern activation as recorded in the intrinsic excitability and synaptic connections of selected neurons (cf. Section 2.5). The localist restriction of parameter updates is the decisive difference to a distributed network update rule, and ensures its specific recall properties. It is also computationally very efficient.

### 3.4 Classification of recalled patterns

The AutoGluon classifier had been trained with the neural representations recorded over 300ms of the 50 variants for each digit, presented to the network before plasticity is applied.

The trained classifier was subjected to the recalled representations on the network, after plasticity has been applied (Table 2, Figure 7). The representations unfold from selected stimulation (*m* = 5, 10, 20) of high-MI neurons. The results are shown in Figure 7. They show, without further analysis, that recall of a complete representation from a few ‘concept’ neurons is possible and that the recalled representations are similar enough to the original representations to be recognized.

**Fig 7.**
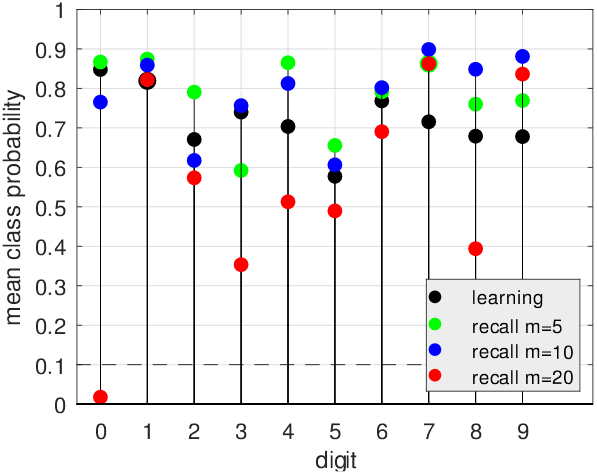
Pattern recall: classification for each digit pattern (**a**) by supervised learning (black) and (**b**) from recalled representations with *m* = 5 (green), *m* = 10 (blue), and *m* = 20 (red). The dashed line shows the chance level *p* = 0.1.

**Fig 8.**
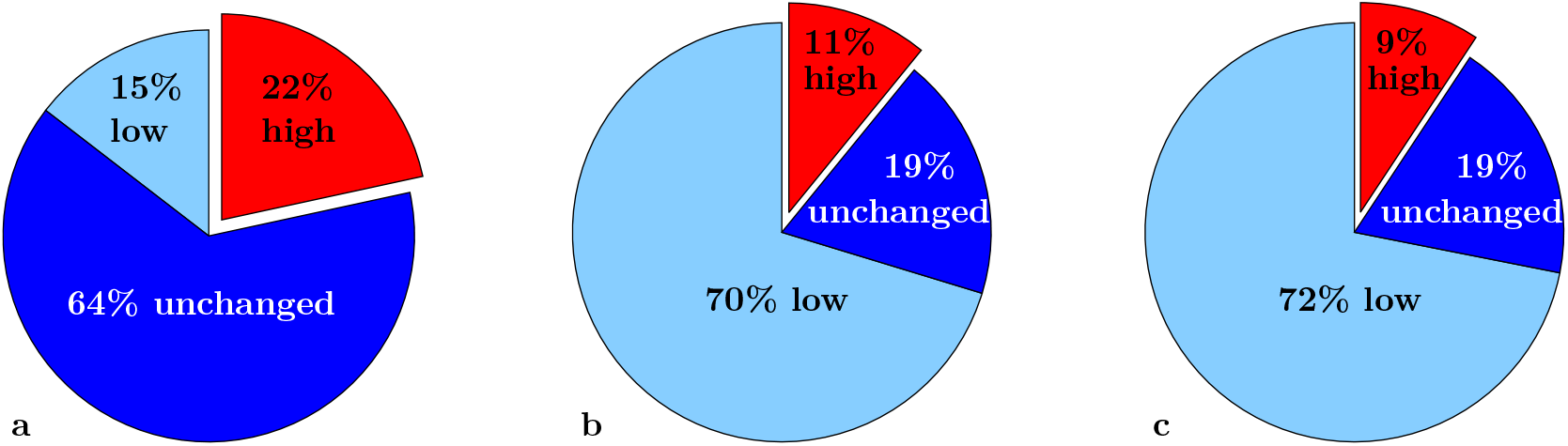
Exemplification of localist learning (mean of 50 variations for “1”): (**a**) patterns prior to plasticity, (**b**) after plasticity, and (**c**) after targeted stimulation with (*m* = 10 per digit). The percentage of E neurons with no response (**a**) becomes low activity in (**b, c**). Number of high-activity neuron (**a**) becomes compressed (**b, c**).

The analysis of errors showed an interesting result: the input representations both before and after plasticity yielded a number of confusions involving the digits, mostly for the patterns “2” and “5”. But the stimulated representations show an error exclusively for “0”, which was always correctly classified before.

What is going on? The structural remap of the pattern representations (cf. Fig. 9) leads to new results similarity and overlap. The input similarity or likelihood of confusion, while kept during plasticity, is not carried over to the stimulated representations. We have arrived at a new ‘symbolic’ transform, the properties of which depend on various factors: the choice of *m*, the selection of inhibitory neurons, the network initialization, etc. It is worthwhile to mention that the stimulated ‘symbolic transform’ is fully deterministic (as long as the background input is non-stochastic). The advantages of a deterministic response to stimulation may become apparent in later applications.

**Fig 9.**
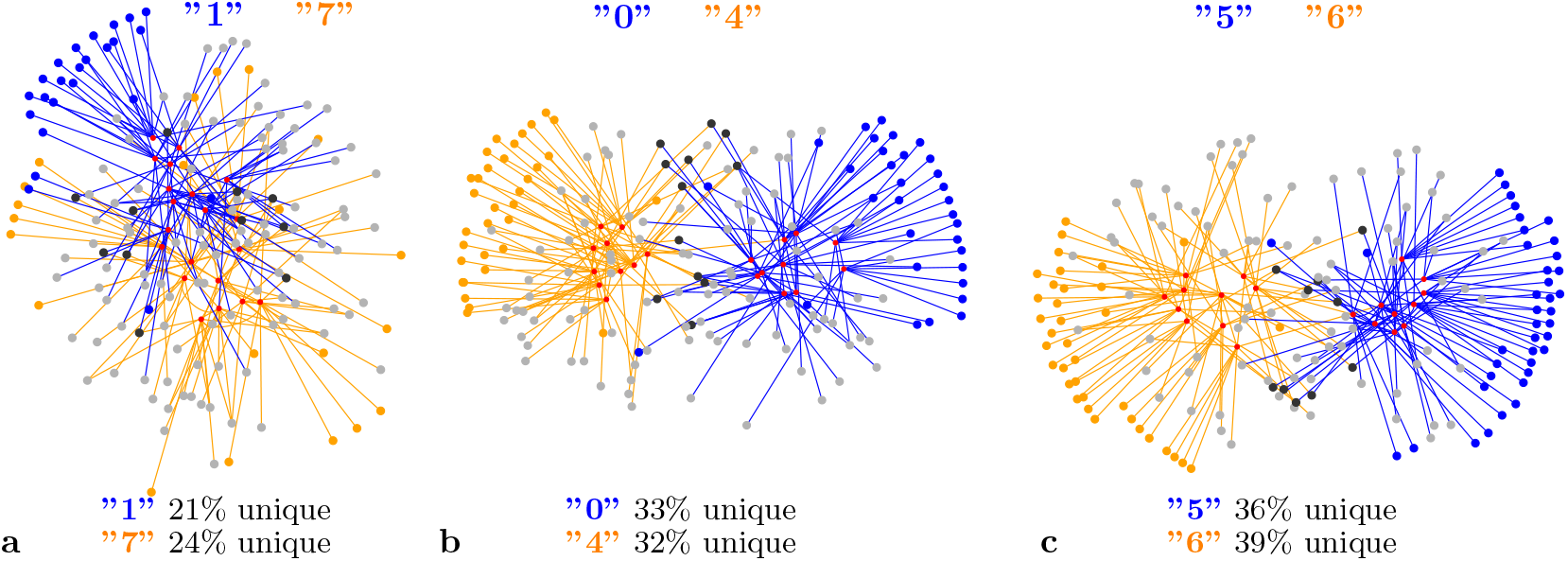
Network connection graph after plasticity for three examples. Target neurons for the strongest 100 connections from high-MI neurons (red) and their connections are shown. Unique feature neurons are colored, black indicates a shared feature neuron, grey feature neurons overlap with other digits. **a**: fewer unique neurons indicating higher similarity, **b**: more unique, and **c**: most unique features.

Nonetheless, we can retrieve a version of the original input representation after plasticity learning. Thus there is local storage at a small set of neurons for large patterns of hundreds of neurons, such that stimulation of these neurons allows to ‘unfold’ and spread activations to retrieve large representations. For each pattern, only a small subset of neurons and synapses are affected by plasticity. As we can see from Fig. 8, there is reduction of activation on all neurons not affected by an input pattern as in Fig. 8**a** (pre-plasticity) vs. **b** (post-plasticity). These percentages are very stable across patterns.

What the percentages show is that in a naive network there is a large number of neurons with minimal response to patterns. In a trained network most neurons show suppressed activation in response to a pattern. about 20% have no response, and about 10% raise their activation level. The naive state could be said to have a ‘reservoir’ of unallocated neurons (Jaeger, 2002; Maass et al., 2002). After plasticity, the number of high activation neurons is somewhat reduced—one could speak of a ‘compressed’ active representation.

### 3.5 Analysis of the results

The goal of the localist plasticity is to allow restoration of representations with high accuracy from activation of high MI neurons alone.

With our model, pattern recall by localist stimulation of such ‘concept’ neurons can be investigated. To understand the process we look at the neural representation after plasticity has been applied. The feature representations for each concept may be overlapping, and the same features can be re-used in multiple representations (cf. Fig.9). Nonetheless, the digit patterns retain a significant proportion of features, which are unique to each digit.

We can also show that the mutual information values for each pattern as the sum of the MI values over the high-ranking neurons increases as a result of plasticity (Fig. 6).

Interestingly, our localist plasticity changes the initial Gaussian distribution to a wider lognormal distribution with heavy-tail characteristics (cf. Figure 10), (Scheler, 2017). The overall distribution of intrinsic gain parameters *a* and *b* after plasticity approaches a lognormal distribution, where the high-MI neurons are substantially strengthened (Figure 10**a**). The picture is more complex for synaptic weights, because only few weights are changed by plasticity in our example. For *m* = 10 high MI (=concept) neurons, only ≈6400 AMPA connections (out of ≈10^6^, that is 0.5%) are adapted. This leads to an uneven distribution of synaptic weights (Figure 10**b**). Extrapolating for larger numbers of weight updates could lead to a biologically attested lognormal distribution.

**Fig 10.**
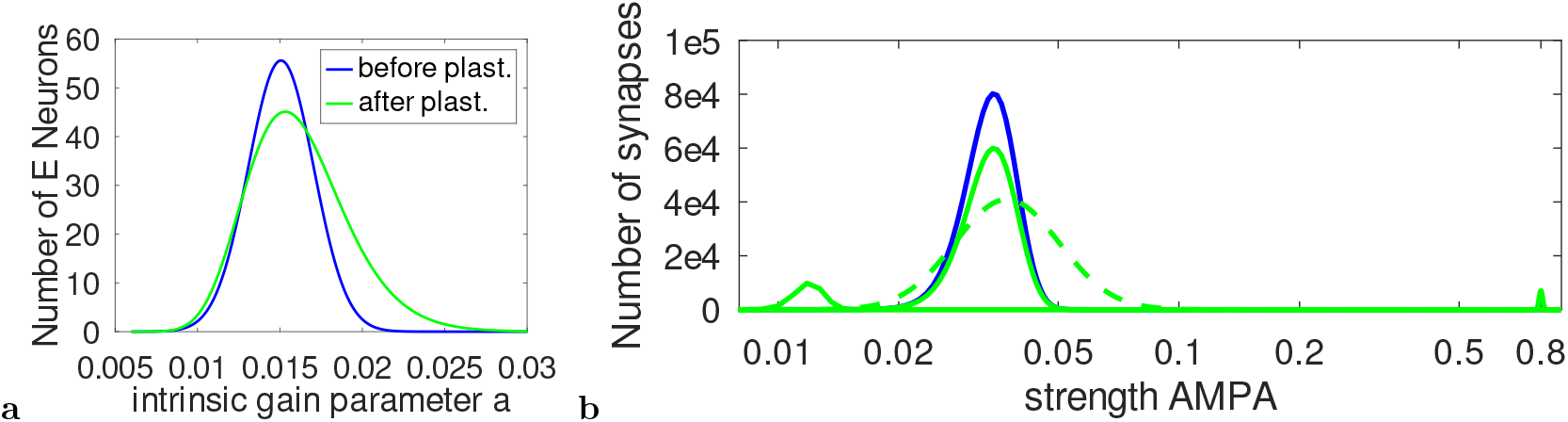
(**a**) Distribution of the intrinsic gain parameters *a* and *b* before applying plasticity (blue), and after plasticity (green). The Gaussian distribution is stretched to a lognormal distribution. (**b**) Distribution of strength of E-E AMPA connections prior to applying plasticity (blue), fitted to a Gaussian. After plasticity, the distribution consists of three different groups (green): strong synapses, which have been strengthened during plasticity, a group of weakened synapses, and a majority of unchanged synapses. We assume that ‘housekeeping’ by homeostatic plasticity (sleep) would lead to an attested distribution.

It is remarkable that all pattern memories are stored in parallel, and stimulation of pattern-specific neurons will reproduce the associated distributed pattern with considerable accuracy over the whole network. Learning a pattern with localist adaptation minimally affects the response of the network to other patterns. The network shows considerable independence in its response to input. Our simple learning rule could be further developed.

The questions of unique features for patterns vs. overlap of features (Fig. 9 colored vs. grey/black target neurons) is important for abstraction and generalization. It is an open questions, which patterns are suitable for a concept-feature ‘symbolic’ abstraction and whether a learning mechanism is successful which *imposes* a structure onto the material for ease of simple one-shot learning. But it is entirely possible that cortical plasticity structures the learning material. This could be complemented by other types of structural learning in cortical and subcortical areas. The enormous simplicity of the plasticity rule and the relative success of the given example problem is to be understood on the background of imposed structure learning.

There are implications from this work for biology and for technical applications.

- biology: We show that a cortical model is able to form high-level ‘concept’ neurons where individual pattern information is stored in a localized manner. A version of the pattern information can be retrieved from these neurons, forming a feature ensemble as in Fig. 9. These results fit with recent experimental work (Josselyn & Tonegawa, 2020). It has been empirically shown that up-regulation of intrinsic excitability occurs for those neurons which are part of an ensemble (engram) (Pignatelli et al., 2018).
- technical: Input patterns are learned as high information ‘concept’ neurons and their connections in a single trial. Over time, a number of different patterns can be stored in this way. Questions of pattern storage and separation, overlap and similarity (generalization, abstraction) by feature neurons are an obvious next step to make technical applications feasible.

## 4 Discussion

We developed a mechanism to store patterns in cortical-like networks using a neuronal ensemble, more precisely a concept-feature ensemble. The plasticity rule stores patterns on the network by localist adaptation of high MI neurons and their synaptic connections, creating a hub-spoke structure. By stimulating only single-pattern high MI ‘concept’ neurons, the resulting representation unfolds on the network and is similar enough to be recognized by a trained classifier. Activating select neurons by direct stimulation results in a whole set of related neurons to recreate a similar pattern to the original. In our network model individual neurons are recruited as pattern storage elements. We thus achieve a localist memory with a distributed component.

Such a concept-feature ensemble need not be just a passive storage of pattern information. The ‘concept’ neurons may also act as ‘control’ neurons, when they are interacting with each other, forming a set of high-level neurons with access to their feature neurons as needed.

It is noticeable that the learning method imposes a structure on the network that is characteristic, could be significant for biological cognition, and well-suited for symbolic computation.

We have not discussed the biological mechanisms which could underlie such selective plasticity, but there are possibilities in the cell-internal memory which filters the information by a number of indicators, such as small molecules and proteins (Scheler, 2023).

The possibilities for structure-imposed pattern abstraction learning by cortical networks are much more comprehensive, involving techniques of control, e.g., for inferences.

We have also not yet analyzed the possibilities for blends and interference between patterns which may serve to create higher-order memory. Our method of representing pattern information in a network allows for efficient storage of concept-feature ensembles. This has been exemplified here for the first time.

## Declarations

The work for this paper did not receive any funding. The authors have no competing interests to declare that are relevant to the content of this article.

## Footnotes

1 NMDA has intrinsic delays of about 120ms

## Notes

### Competing Interest Statement

The authors have declared no competing interest.

### Summary of Updates

Text and figures revised and updated

